# *In situ* cell-only bioprinting of patterned prevascular tissue into bioprinted high-density stem cell-laden microgel bioinks for vascularized bone tissue regeneration

**DOI:** 10.1101/2025.03.17.643708

**Authors:** Oju Jeon, Hyoeun Park, Min Suk Lee, Eben Alsberg

**Affiliations:** Departments of Biomedical Engineering, University of Illinois Chicago, IL 60612, USA; Department of Biomedical Engineering, Dongguk University, Goyang, Gyeonggi 10326, Republic of Korea; Departments of Orthopaedic Surgery, University of Illinois Chicago, IL 60612, USA; Departments of Mechanical & Industrial Engineering, University of Illinois Chicago, IL 60612, USA; Departments of Pharmacology and Regenerative Medicine, University of Illinois Chicago, IL 60612, USA; Jesse Brown Veterans Affairs Medical Center (JBVMC), Chicago, IL 60612, USA

## Abstract

Recently, microgels have been widely used in three-dimensional (3D) bioprinting as both supporting baths and bioinks. As a bioink, microgels have several unique properties, such as shear-thinning and self-healing behaviors with tunable mechanics, making them useful in 3D bioprinting. While cell encapsulated microgels offer many advantages in 3D bioprinting, they also have some limitations. It is still challenging to produce large quantities of cell encapsulated microgels with consistent quality and properties due to processes that are often complex and time-consuming. In this study, stem cell encapsulated, photocrosslinkable, shear-thinning and self-healing alginate microgel (SSAM) bioinks have been successfully fabricated via simple mixing of an oxidized and methacrylated alginate solution with suspended stem cells and a supersaturated calcium sulfate slurry solution through a custom-made spiral mixing unit. The SSAM bioinks can be bioprinted into complex 3D structures with both high resolution and shape fidelity due to their shear-thinning and self-healing properties. The 3D bioprinted SSAM bioinks can then serve as a supporting bath for the creation of prevascular network patterns using an individual cell-only prevasculogenic bioink within the 3D printed constructs. The prevascular network patterned 3D bioprinted constructs can be further stabilized by secondary photocrosslinking of the SSAMs, which enables long-term culture of the printed constructs for functional vascularized osteogenic tissue formation by differentiation of the bioprinted cells. The SSAM bioinks and individual cell-only printing technique enable *in situ* bioprinting of prevascularized tissue constructs in a mouse calvarial bone defect, achieving mechanical stability and ensuring the *in situ* bioprinted constructs remain within the defect.

## Introduction

The bioprinting field has revolutionized tissue engineering by enabling the fabrication of complex three-dimensional (3D) tissue constructs that closely mimic the structural and functional characteristics of native tissues [1, 2]. The development of bioinks, using biocompatible materials that support cell viability and allow for precise control over spatial organization during the printing process, is pivotal to these advancements [3, 4]. Among various bioinks, microscale crosslinked hydrogel particles (microgels) have gained great attention due to their versatility and favorable properties in supporting cell encapsulation [5–9]. Microgels can serve as both the bioink itself [8–11] and a supportive medium [8], providing structural integrity and a hospitable microenvironment for embedded cells [12, 13]. Particularly, living cell-encapsulated microgel bioinks formed with biodegradable materials are valuable in tissue engineering applications, as they enable high resolution and shape fidelity 3D printing with high cell viability and have tunable degradation rates [10, 14, 15].

Despite these advantages, the use of cell encapsulated microgel bioinks in tissue engineering faces several notable challenges. The production processes for these bioinks are often complex and require precise control over various parameters to maintain cell viability during encapsulation [16–18]. Scalability in sterile environments is also challenging as microgel production methods must be adapted to meet clinical standards while maintaining sterility [12, 19–21]. Furthermore, some bioink preparation processes involve potentially toxic reagents, which can compromise cell health and limit the bioink’s clinical applicability [17, 22]. Addressing these limitations is essential for the practical use of microgel bioinks in tissue engineering and regenerative medicine. In this study, to address these challenges, we developed a robust, scalable, and cytocompatible method for producing a stem cell-laden, photocrosslinkable, shear-thinning, and self-healing alginate microgel (SSAM) bioink. This was achieved by simply mixing human mesenchymal stem cells (hMSCs) suspended in an oxidized and methacrylated alginate (OMA) solution with a supersaturated calcium sulfate slurry solution using a custom-designed spiral mixing unit. Our approach maintained sterility throughout the process and allowed for efficient, reproducible fabrication of hMSC-laden microgels, addressing key challenges in bioink formulation and bioprinting applications.

Vascularization is essential for engineering clinically relevant tissue constructs, as it facilitates oxygen and nutrient supply, waste removal, and integration with host tissue post-implantation [23]. Numerous studies have highlighted the importance of prevascularization in engineered tissue constructs, demonstrating that preformed prevascular networks significantly enhance tissue survival, function and long-term viability after implantation [23–27]. Prevascular networks can significantly improve perfusion and nutrient exchange within engineered tissue constructs, allowing them to better mimic the complex physiological microenvironment of native tissues [28, 29]. In this study, by bioprinting patterned individual cell-only prevasculogenic bioinks into our constructs, we aim to further advance the formation of prevascular structures that can potentially enhance osteogenic differentiation of hMSCs and promote overall osteogenic tissue formation.

This approach has the potential to pave the way for the development of functional, vascularized tissues that meet the complex demands of clinical applications. Ultimately, it could advance the field of bioprinting and significantly impact regenerative therapies by enabling the fabrication of highly functional tissue constructs suitable for a wide range of clinical needs.

## Materials and Methods

### Synthesis of OMAs

Oxidized alginates (OAs) were prepared by reacting sodium alginates with sodium periodate. Sodium alginate (10g, Protanal LF120M, 251 mPa•S at 1% w/v in water at 25°C, IFF) was dissolved in ultrapure deionized water (diH_2_O, 900ml). Sodium periodate (0.218 and 0.545g, Sigma) was dissolved in diH_2_O (100ml), added to sodium alginate solution under vigorous stirring to achieve 2 and 5% theoretical alginate oxidation, respectively, and allowed to react in the dark at room temperature for 24 hours. Methacrylation (20% theoretical) was performed to obtain OMAs by reacting OAs with 2-aminoethyl methacrylate hydrochloride (AEMA, Polysciences). To synthesize OMAs, 2-morpholinoethanesulfonic acid (MES, 19.52g, Sigma) and sodium chloride (17.53g, Fisher) were directly added to each OA solution (1L) and then the pH was adjusted to 6.5. N-hydroxysuccinimide (NHS, 1.176g, Sigma) and 1-ethyl-3-(3-dimethylaminopropyl)-carbodiimide hydrochloride (EDC, 3.888g, Oakwood Chemical) were added to the mixtures under stirring to activate 20 % of the carboxylic acid groups of the sodium alginate. After 5 minutes, AEMA (1.688g) was added to each solution, and the reactions were maintained in the dark at room temperature for 24 hours. The reaction mixtures were precipitated into excess acetone and dried in a fume hood. After rehydrating to an 1% w/v solution in diH_2_O, the OMAs were purified by dialysis against diH_2_O using a dialysis membrane (MWCO 3500, Spectrum Laboratories) for 3 days, treated with activated charcoal (5g/L, 100 mesh, Oakwood Chemical) for 30 minutes, filtered (0.22μm filter, Millipore Sigma) and lyophilized. To determine the levels of alginate methacrylation, the OMAs were dissolved in deuterium oxide (D_2_O, 2% w/v, Sigma), and ^1^H-NMR spectra were recorded on an NMR spectrometer (600 MHz, Bruker) using 3-(trimethylsilyl)propionic acid-d_4_ sodium salt (0.05% w/v, Sigma) as an internal standard [30–32].

### Fabrication of the SSAM bioinks

Bone marrow-derived hMSCs from a single donor were isolated using a Percoll^®^ gradient (Sigma) and the differential adhesion method [33, 34]. Briefly, bone marrow aspirate was obtained from the posterior iliac crest of a donor under a protocol approved by the University Hospital of Cleveland Institutional Review Board. The bone marrow aspirate was washed with DMEM-LG (Sigma) containing 10% prescreened fetal bovine serum (FBS, Sigma), 100U/mL penicillin and 100μg/mL streptomycin (1% P/S, BioWhittaker). Mononuclear cells were isolated by centrifugation in a Percoll^®^ density gradient, and then isolated mononuclear cells were plated at 1.8×10^5^ cells/cm^2^ in DMEM-LG containing 10% FBS and 1% P/S. After 4 days of culture in an incubator at 37°C and 5% CO_2_, non-adherent cells were removed, and adherent cells were maintained in DMEM-LG containing 10% FBS and 1% P/S with media changes every 3 days. After 14 days of culture, the cells were passaged at a density of 5 × 10^3^ cells/cm^2^, cultured for an additional 14 days, and then stored in cryopreservation media (90% FBS and 10% DMSO) in liquid nitrogen until use.

To fabricate the SSAM bioinks, OMAs (100mg) were dissolved in DMEM-LG (5ml) containing 0.05 % photoinitiator [2-Hydroxy-4’-(2-hydroxyethoxy)-2-methylpropiophenone, Sigma]. hMSCs were expanded in a media consisting of DMEM-LG with 10% FBS and 1 % P/S and harvested with trypsin/EDTA (Thermo Fisher) and concentrated by centrifugation at 300×g for 5 min. Following aspiration of the supernatant, pelleted hMSCs (passage number 4) were suspended in OMA solutions (1 × 10^7^ cells/ml), and then the OMA solutions with suspended hMSCs (5 ml) were placed in a 10 ml syringe. 200µl of supersaturated CaSO_4_ (0.21 g/ml in diH_2_O) was loaded into another 10 ml syringe. After the two syringes were connected with a custom-made female-female lure lock spiral mixing unit (**Supporting Figure 1**), the two solutions were mixed back and forth 40 times, and then they were further mixed back and forth 10 times every 10 minutes for 30 minutes. Viability of hMSCs in the SSAM bioinks was evaluated using Live/Dead staining comprised of fluorescein diacetate (Sigma) and ethidium bromide (Fisher) [8, 10]. After suspending the SSAM bioinks (100 μl) in 1 ml of expansion media, 20 μl of staining solution was added into the suspension and incubated for 3-5 minutes at room temperature, and then stained SSAM bioinks were imaged using a fluorescence microscope (ECLIPSE TE300, Nikon) equipped with a digital camera (AmScope). To visualize the hMSC encapsulated SSAMs, they were stained with Safranin O (1%, Fisher) and then imaged.

### Rheological properties of the SSAM bioinks

Rheological properties of the SSAM bioinks were measured with a rheometer (Kinexus Ultra+, Malvern Panalytical) to evaluate their shear-thinning behavior, shear-yielding at low shear stress and self-healing properties. In oscillatory mode, a parallel plate (8mm diameter) geometry measuring system was employed, and the gap was set to 1 mm. After each hMSC encapsulated SSAM bioink was placed between the plates, all the tests were started at 25 ± 0.05°C, and the plate temperature was maintained at 25°C. To determine the shear-thinning behavior and shear-yielding at low shear stress of the SSAM bioinks, viscosity changes were measured as a function of shear rate and shear stress, respectively. Oscillatory frequency sweep (0.1-10 Hz at 1 % shear strain) tests were performed to measure storage modulus (G′) and loss modulus (G″). Oscillatory strain sweep (0.10-100 % shear strain at 1 Hz frequency) tests were performed to determine the G’/G” crossover. To demonstrate the self-healing properties of the hMSC encapsulated SSAM bioinks, cyclic deformation tests were performed at 100 % shear strain with recovery at 1 % shear strain, each at 1 Hz for 1 minute.

### Resolution analysis of 3D printed the SSAM bioinks

The SSAM bioinks were loaded into 3ml syringes (BD), connected to 0.5 inch printing needles, and mounted on a commercial syringe pump-based 3D bioprinter (BioX, Cellink). The tip of each needle was positioned on the bottom of a petri dish, and the print instructions were sent to the printer using the in-house software in the printer. Linear SSAM bioink filaments were printed with 27, 25, 23, 22, 20 and 18G printing needles and then imaged using a microscope (ECLIPSE TE300) equipped with a digital camera (MU Series, AmScope). The diameters of the 3D printed hMSC encapsulated SSAM bioink filaments were measured at 10 different locations on 5 filaments for each bioink using ImageJ (National Institutes of Health).

### Preparation of individual cell-only prevasculogenic bioink

Human adipose tissue-derived stromal cells (hASCs) were generously provided by the Stem Cell Biology Laboratory at the Pennington Biomedical Research Center (Baton Rouge, LA) [35]. Human umbilical vein endothelial cells (HUVECs) were purchased from ATCC. The HUVECs were expanded in endothelial cell growth medium-2 (EGM-2, Lonza). hASCs (passage 3) and HUVECs (passage 4) (1:1 ratio) were mixed to prepare individual cell-only prevasculogenic bioinks.

### 3D bioprinting of prevascular network patterned 3D constructs

The SSAM bioinks were prepared as described above with the following modification. To functionalize the SSAM bioinks with RGD peptide for promoting interactions with the individual cell-only prevasculogenic bioinks, RGD peptides (2 mg, CGGGRGDSP, GenScript) were dissolved with OMA (100mg) in DMEM-LG (5 ml) containing 0.05 % photoinitiator. After 3D printing SSAM bioinks with 22G or 20G printing needles using the BioX printer (10×10×3 mm cuboids), the individual cell-only prevasculogenic bioinks were used to 3D print 2 grid patterns in the z-plane with a 25G printing needle into the 3D SSAM bioink printed constructs with 1 mm spaces in all directions between printed filaments. For comparison, a mixture of the RGD-modified SSAM bioinks and the individual cell-only prevasculogenic bioinks were prepared at the same volume ratio (SSAM bioink:individual cell-only prevasculogenic bioink = 50:3) as the patterned construct printing and then 3D printed. As a control group, constructs using only the RGD-modified SSAM bioink were also 3D printed. After photocrosslinking under UV light at 20 mW/cm^2^ for 1 minute, 3D bioprinted prevascular network patterned constructs were culture in mixed media [2×concentrated osteogenic media (DMEM-HG containing 200nM dexamethasone (Sigma), 75μg/ml L-ascorbic acid-2-phosphate (Wako), 20mM beta-glycerophosphate (Sigma), 20% FBS, 2% PS, and 200ng/ml BMP-2 (GenScript)):2×concentrated EGM-2 = 1:1]. After 4 weeks of culture, 3D printed prevascular network patterned constructs were fixed in 10 % neutral buffered formalin overnight at 4 °C, embedded in paraffin, sectioned at a thickness of 10 µm, stained with Alizarin red S and CD31, and then imaged using a using a microscope (ECLIPSE TE300) equipped with a digital camera (MU Series). For quantification of alkaline phosphatase (ALP) activity, DNA content and calcium deposition, osteogenically differentiated 3D printed prevascular network patterned constructs were homogenized at 35,000 rpm for 60 seconds using a TH homogenizer (Omni International) in 2 ml ALP lysis buffer (CelLytic M, Sigma). The homogenized solutions were centrifuged at 500×g with a centrifuge (Sorvall Legend RT Plus, Thermo Fisher Scientific). For ALP activity measurement, supernatant (100μl) was treated with *p*-nitrophenylphosphate ALP substrate (pNPP, 100 μl, Sigma) at 37 °C for 30 minutes, and then 0.1 N NaOH (50 μl) was added to stop the reaction. The absorbance was measured at 405 nm using a plate reader (Molecular Devices). A standard curve was made using the known concentrations of 4-nitrophenol (Sigma). DNA content in supernatant (100 μl) was measured using a Quant-iT PicoGreen assay kit (Invitrogen) according to the manufacturer’s instructions. After an equal volume of 1.2 N HCl was added into each lysate solution to dissolve the calcium compounds, the mixed solutions were centrifuged at 500×g with a centrifuge (Sorvall Legent RT Plus). The amount of calcium in the supernatant was measured using a calcium assay kit (Pointe Scientific) according to the manufacturer’s instructions. All ALP activity and calcium deposition measurements were normalized to DNA content.

### RNA isolation and real-time quantitative reverse transcription-polymerase chain reaction (qRT-PCR)

RNA was isolated from the osteogenically differentiated 3D printed prevascular network patterned constructs using TRI reagent (Sigma) according to the manufacturer’s instructions, and first-strand cDNAs were synthesized using a cDNA synthesis kit (Gold Biotechnology) according to the manufacturer’s instructions. qRT-PCR was performed with a SYBR®Premix Ex Taq TM kit (Takara Bio) using a real-time thermal cycler (qTOWER^3^ 84G, Analytik Jena) according to the manufacturer’s instructions. The primer sequences used for qRT-PCR are in **Supporting Table 1**. Relative mRNA expression for the target gene of interest (TGI) was normalized to endogenous control GAPDH gene using the delta threshold cycle (ΔCt) method. The Ct for each gene and endogenous GAPDH in each sample was used to create ΔCt_TGI_ values (Ct_TGI_-Ct_GAPDH_). ΔΔCt values were calculated by subtracting the ΔCt_TGI_ of the control (no prevascularization) from the ΔCt_TGI_ of experimental groups. The relative expression of the target genes was then calculated (relative fold gene expression = 2^-ΔΔCt^).

### Evaluating *in situ* bioprintability

An *in situ* printability experiment was performed using a mouse cadaver, obtained from the Biological Research Laboratory Animal Facilities at the University of Illinois Chicago. After shaving the skin, a midline sagittal incision was made to expose the calvarium. The skin and periosteum were elevated and retracted. Two circular, 4-mm diameter defects on the parietal bone, one on each side of the sagittal suture, were created by drilling with a high speed Dremel rotary tool to minimize skull crushing and cracking. The full thickness of the bone was removed, and the defect sites were cleaned with gauze and saline irrigation. After calibrating the location of the printing nozzles to the defect site, the SSAM bioink was *in situ* 3D bioprinted using a 22G printing needle into a calvarial bone defect (4 mm diameter and 0.5 mm thickness). Next, the individual cell-only prevasculogenic bioink was 3D printed in a grid pattern using a 25G printing needle with 1 mm spaces between printed filaments within the *in situ* 3D bioprinted SSAM slurry structure. After photocrosslinking the 3D printed construct under UV light at 20 mW/cm^2^ for 1 min, the photocrosslinked construct was retrieved.

### Statistical analysis

Statistical analyses were performed by one-way analysis of variance (ANOVA) with the Tukey significant difference post hoc test using the Prism software (Graphpad). A value of *p*<0.05 was considered statistically significant.

## Results and Discussion

Microgels, microscale crosslinked hydrogel particles, are widely used in bioprinting application as bioinks and as a supporting bath for embedding 3D printing [8–10, 14, 15]. Living cell encapsulated microgel bioinks in particular are often used as building blocks for scaffolding in tissue engineering [10, 14]. While sometimes cell encapsulated microgel bioinks offer advantages over traditional bulk hydrogel bioinks for tissue engineering, such as enhanced cell viability during printing, high resolution printing, and/or improved diffusion of nutrients, oxygen and waste products, and structural integrity, there are still several limitations that hinder their application, including complicated production processes that may also be toxic to cells and scaling up fabrication while maintaining sterile conditions [15, 19]. Here, ionically crosslinked SSAM bioinks have been successfully fabricated via the simple mixing of an hMSCs-suspended OMA solution with a calcium sulfate slurry using a custom-made spiral mixing unit (**Figure 1A**). The 3D printed SSAM bioink can then serve as a supporting bath for the creation of a prevascular network pattern using an individual cell-only prevasculogenic bioink, comprised of HUVECs and hASCs (**Figure 1B**). The 3D bioprinted individual cell-only prevasculogenic bioink pattern and entire 3D printed construct can be further stabilized by secondary photocrosslinking, enabling long-term culture of 3D printed constructs for the formation of vascularized functional tissue by the differentiation of encapsulated hMSCs with an incorporated spatially patterned prevascular network of HUVECs/hASCs condensations.

**Figure 1.**
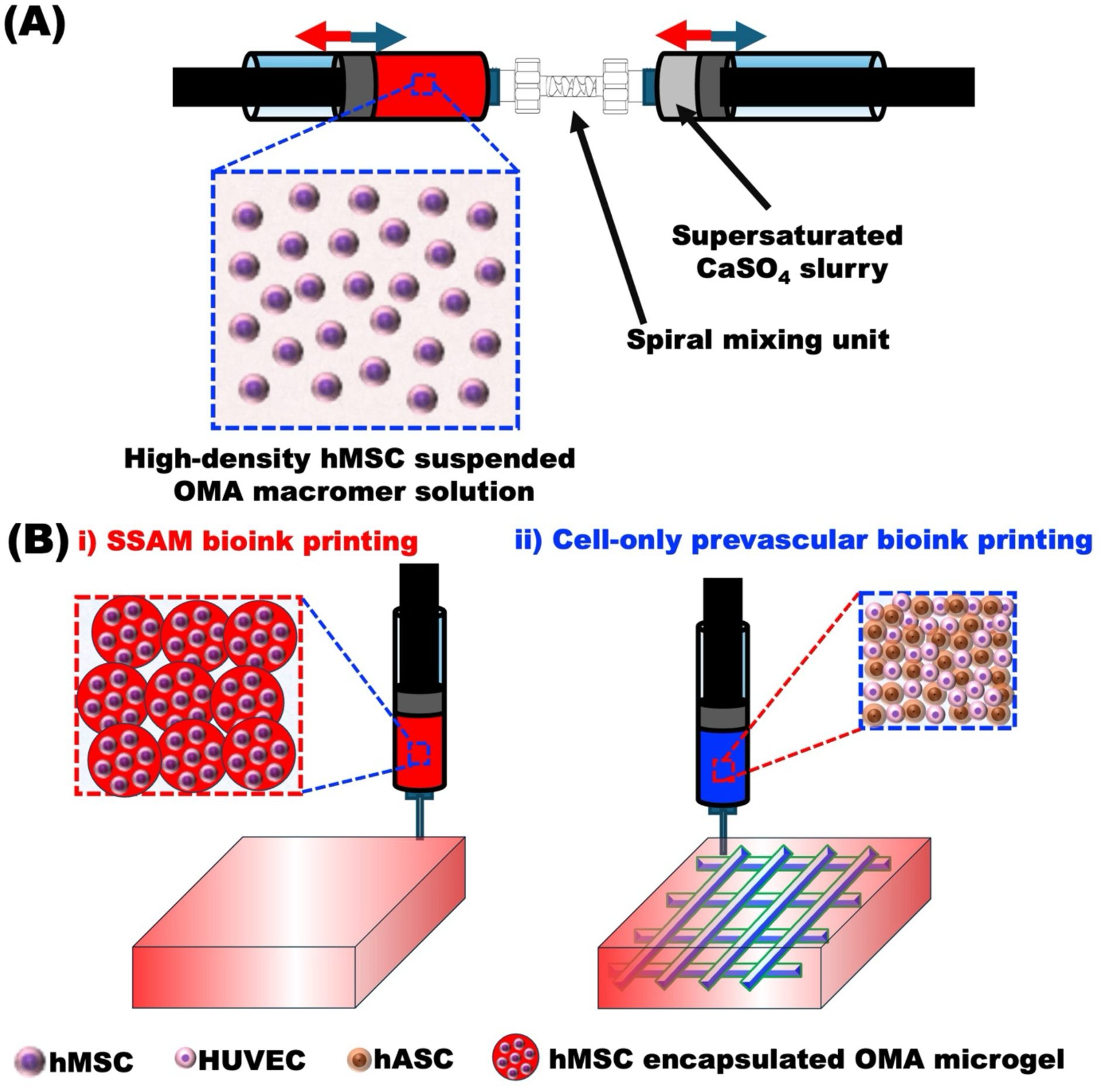
**(A)** Schematic illustration of one-step preparation of hMSC encapsulated SSAM bioink. **(B)** Schematic illustration of 3D bioprinting of the **i)** SSAM bioink to serve as a supporting bath for bioprinting of **ii)** patterned cell-only prevasculogenic bioink to produce a prevascular network patterned construct.

Since cells are exposed to shear forces and hydrostatic and osmotic pressure during the encapsulation process, achieving high cell viability can be challenging [36, 37]. However, in this study, high cell viability was observed after cell encapsulation (**Figure 2A**), indicating that the OMA macromer solution, ionic crosslinking and microgel fabrication processes utilized by our custom-made spiral mixing unit were non-toxic to the encapsulated cells. The SSAM bioinks fabricated using our custom-made spiral mixing unit with 2OX20MA and 5OX20MA OMAs exhibited similar viability (**Figure 2A**), morphology and mean diameters (**Figure 2B**). These findings indicate that our cell-laden microgel fabrication method allows for reliable and cytocompatible cell encapsulation, which is crucial for applications in tissue engineering and regenerative medicine using cell-laden microgels.

**Figure 2.**
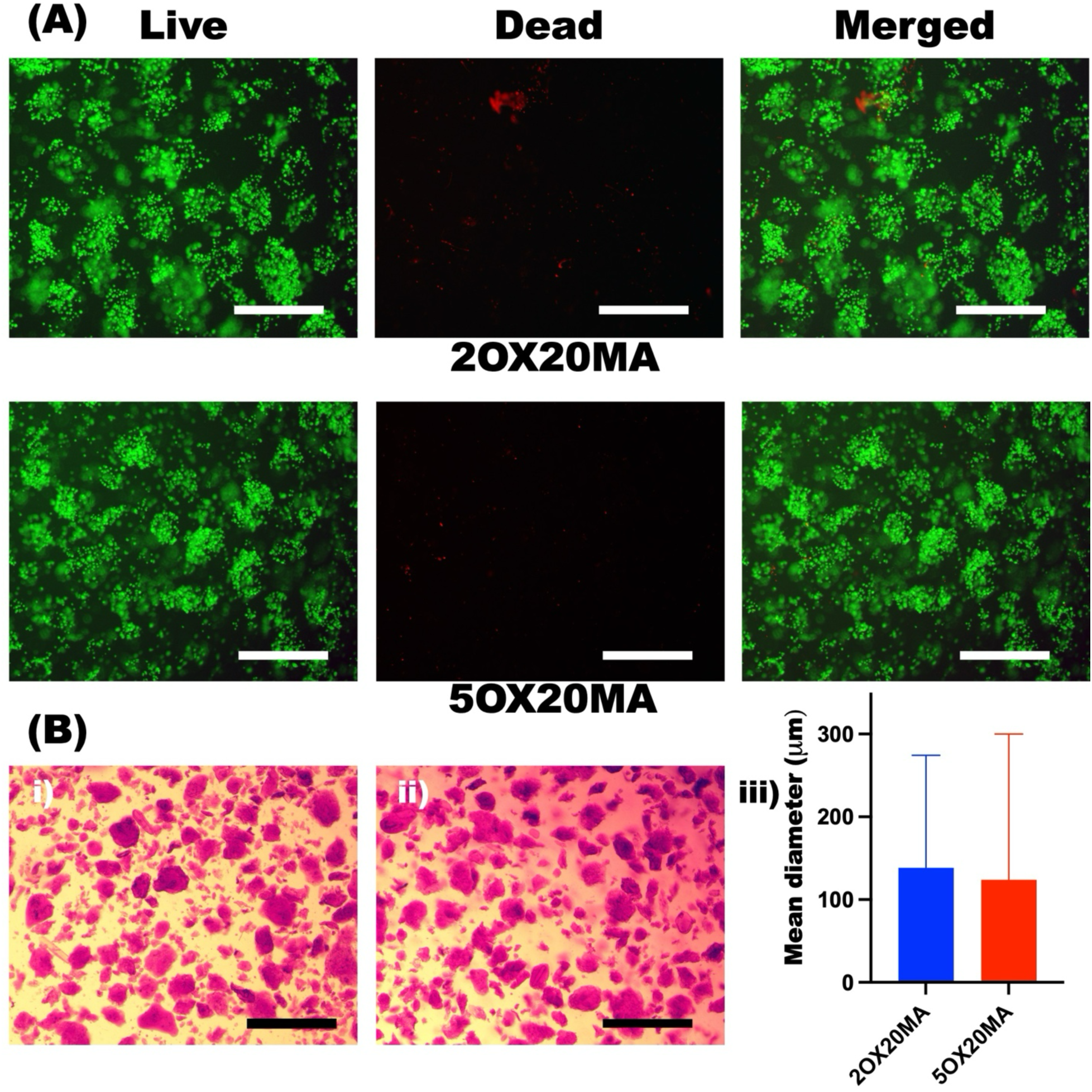
**(A)** Representative Live, Dead and Merged photomicrographs of the SSAM bioinks formed with 2OX20MA and 5OX20MA OMAs. Scale bars indicate 250 μm. **(B)** Photomicrographs of the Safranin O stained SSAM bioinks formed with **i)** 2OX20MA and **ii)** 5OX20MA, and **iii)** quantification of their mean diameters (N=240). Scale bars indicate 500 μm.

Bioprinting is a powerful tool for creating complex 3D tissue structures for tissue engineering and regenerative medicine [1]. For successful 3D bioprinting, bioinks must meet specific key requirements to ensure their printability to precisely create complex 3D structures and accurately form desired shapes. Shear-thinning, shear-yielding, and self-healing properties are essential for bioinks used in 3D bioprinting [16]. Low shear-yielding, shear-thinning and transition from solid-like to liquid-like behavior under shear stress allow bioinks to flow easily under low shear force during extrusion through a narrow printing needle, while self-healing from liquid-like to solid-like enables rapid recovery of the mechanical properties of the bioinks, preserving the integrity of the printed structures and their shapes [16]. Therefore, we evaluated whether the SSAM bioinks exhibit these properties before using them for bioprinting. The SSAM bioinks exhibited shear-thinning behavior, as indicated by a decrease in viscosity with increasing shear rate (**Figure 3A**). Additionally, shear stress sweep tests showed that the SSAM bioinks had a low shear yield stress (**Figure 3B**, yield stress < 2 Pa), indicating that only a small amount of stress is needed for the SSAM bioinks to flow. Oscillatory frequency sweet tests with the SSAM bioinks showed that G’ (storage modulus) was significantly higher than G” (loss modulus), indicating that the SSAM bioinks were mechanically stable (**Figure 3C**). In oscillatory shear strain sweep tests, G” surpassed G’ at approximately 50% shear strain (**Figure 3D**), indicating a phase transition from solid-like to liquid-like, which facilitates the extrusion of the SSAM bioinks. Furthermore, the SSAM bioinks exhibited self-healing characteristics when investigated under cyclic shear strain sweep tests by alternating between 1% and 100% shear strain. The SSAM bioinks showed consistent responses of shear moduli crossover (G”>G’) to 100% (high) shear strain and rapid and repeatable recoveries to their mechanically stable state (G’>G”) at 1% (low) shear strain (**Figure 3E**). Additional subsequent photocrosslinking could enhance the mechanical properties of the SSAM bioinks (**Figure 3F**), enabling long-term culture of 3D bioprinted constructs for functional tissue formation through differentiation of encapsulated hMSCs. Similar to the 2OX20MA SSAM bioinks, the 5OX20MA SSAM bioinks were mechanically stable and exhibited shear-thinning behavior, low shear yield stress, phase change at approximately 90 % shear strain, and self-healing properties (**Supporting Figure 2**).

**Figure 3.**
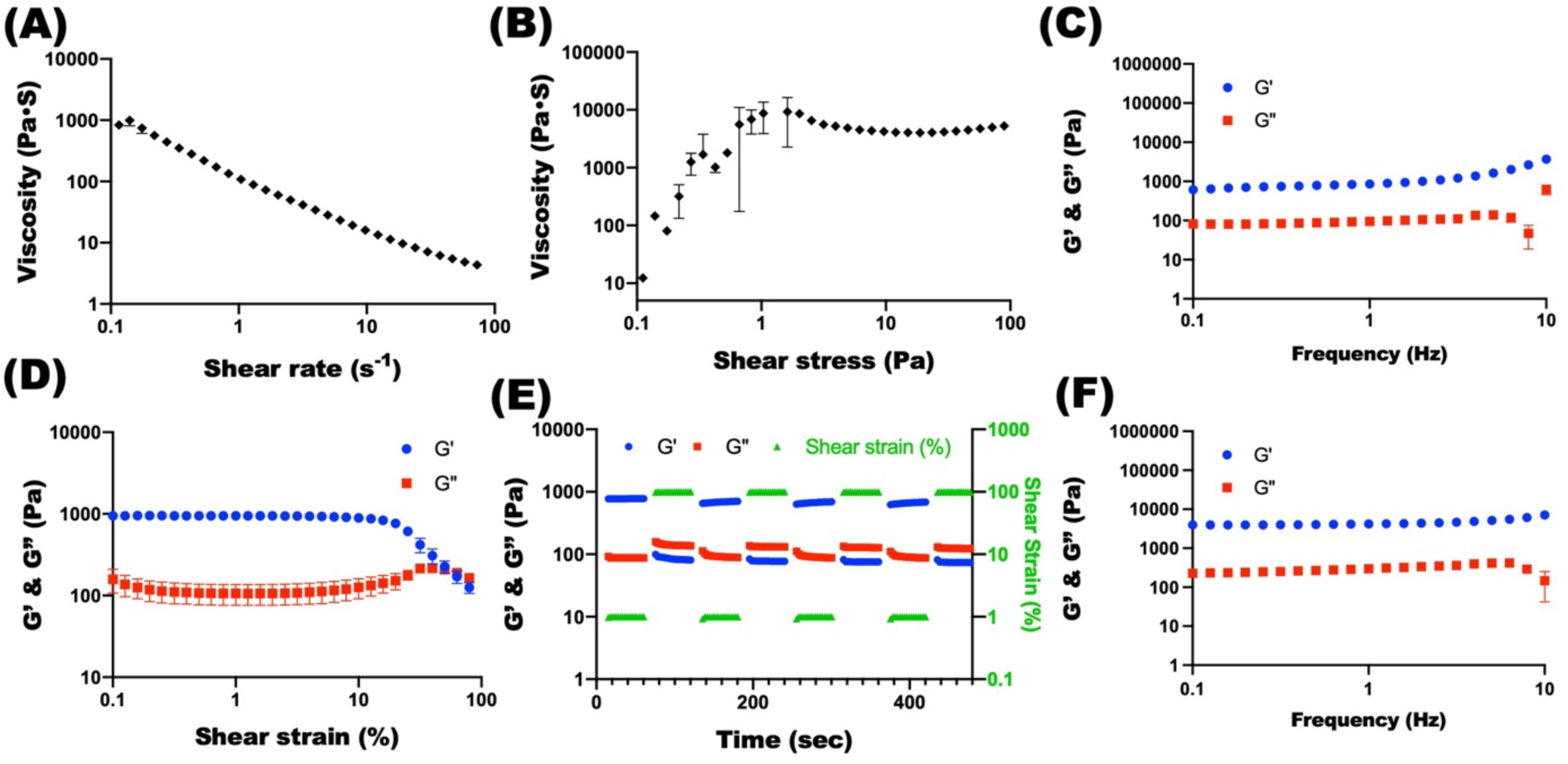
Viscosity measurements of the hMSC encapsulated SSAM bioinks as a function of **(A)** shear rate and **(B)** shear stress demonstrate their shear-thinning and shear-yielding behaviors (N=3). **(C)** Frequency sweep tests indicate that the hMSC encapsulated SSAM bioinks were mechanically stable (N=3). **(D)** Strain sweep tests of the hMSC encapsulated SSAM bioinks indicate their gel-sol transition at approximately 50 % shear strain (G’ and G” crossover) (N=3). **(E)** Shear moduli changes during dynamic shear strain test of the hMSC encapsulated SSAM bioinks with alternating low (1%) and high (100%) shear strains at 1 Hz frequency demonstrate their self-healing or thixotropic property by showing rapid transition between solid-like and liquid-like behaviors. **(F)** Frequency sweep tests of the photocrosslinked hMSC encapsulated SSAM bioinks indicate that additional photocrosslinking enhanced their mechanical strength (N=3).

For precisely mimicking the architectures of native tissues using extrusion-based 3D bioprinting techniques, it is essential to evaluate the resolution of the bioinks [16]. The size of printing nozzles plays a pivot role in determining the resolution of the 3D printed constructs, directly impacting the precision and fidelity of the bioprinted structures [38]. Therefore, the effect of the printing needle diameters on the filament resolution was investigated. To demonstrate the robustness of high resolution printing using the SSAM bioinks, the diameters of the printed filaments were measured using various printing needle gauges from 27G to 18G (**Figure 4A**). As the inner diameter of the printing needles increased from 210 μm (27G) to 838 μm (18G), the diameters of the printed hMSC-laden SSAM bioink filaments also significantly increased from 177 μm to 801 μm (**Figure 4B**). These values ranged from 84 to 114% of the inner diameters of the printing needles, confirming the capacity of our SSAM bioinks to be used for high resolution bioprinting.

**Figure 4.**
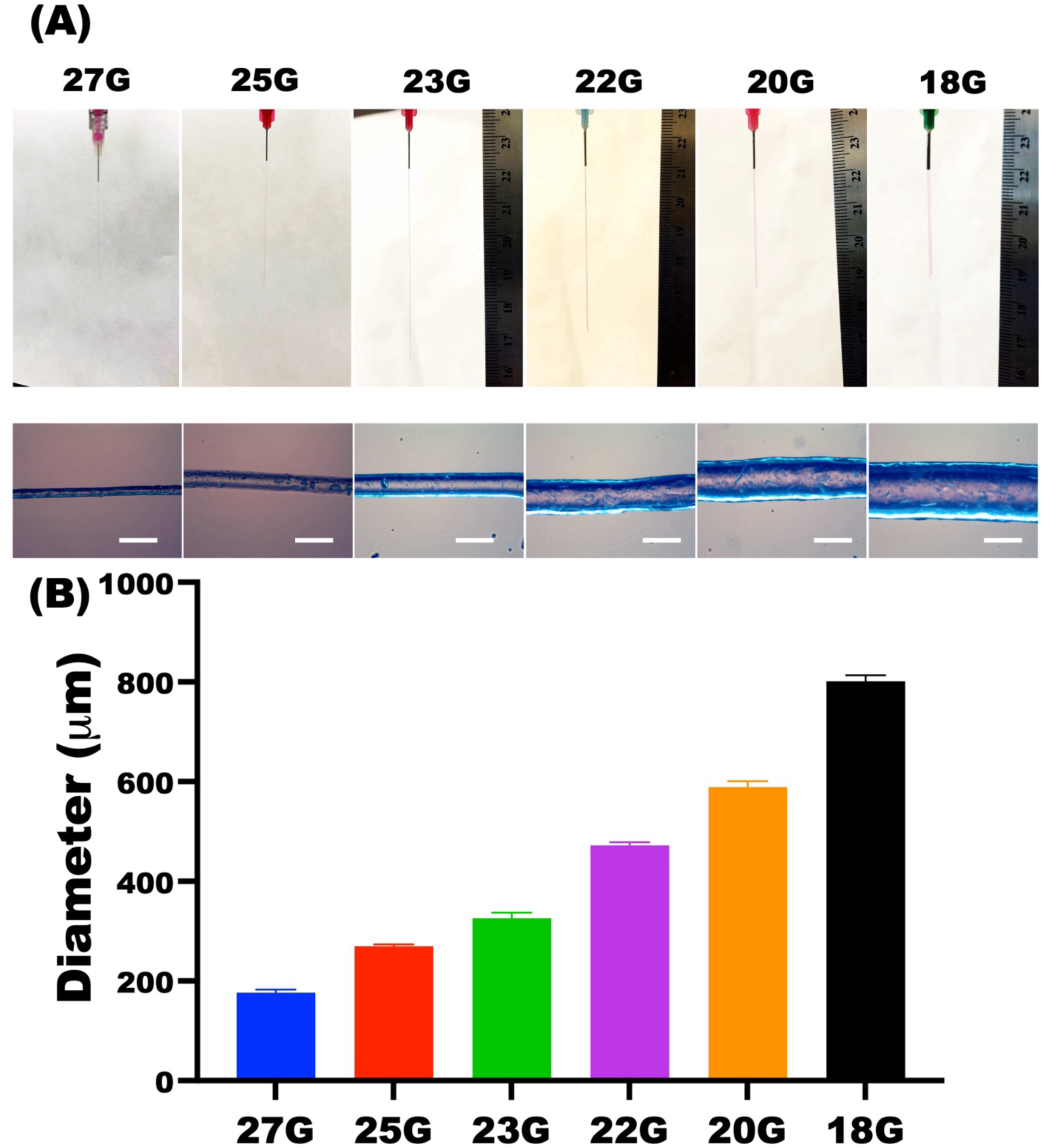
**(A)** Photographs (top) and photomicrographs (bottom) of the extruded hMSC-laden SSAM bioink filaments through various printing needles and **(B)** their mean diameters (N=5).

To demonstrate the ability to use the SSAM bioinks in bioprinting macroscale shapes with high shape fidelity, cuboids (10×10×3 mm) were printed with 22G and 20G printing needles (**Figure 5A**), the dimensions of the 3D sprinted constructs were measured (X, Y and Z axes), and then the shape fidelity as calculated by comparing these dimensions with the 3D digital models. The quantified shape fidelity of the 3D printed structures ranged from 96 to 114 % (**Figure 5B and C**), indicating high shape fidelity. Additionally, there was no significant difference in the shape fidelity between the 22G and 20G printing needles when using 2OX20MA and 5OX20MA SSAM bioinks. These results indicate that the SSAM bioinks enables precise biofabrication of 3D structures, which is crucial for maintaining accurate geometries in tissue engineering. Achieving high shape fidelity in the 3D printed structures regardless of printing needle gauge suggests the versatility of our SSAM bioinks across various extrusion conditions, allowing for flexible integration into a wide range of bioprinting systems without sacrificing high shape fidelity.

**Figure 5.**
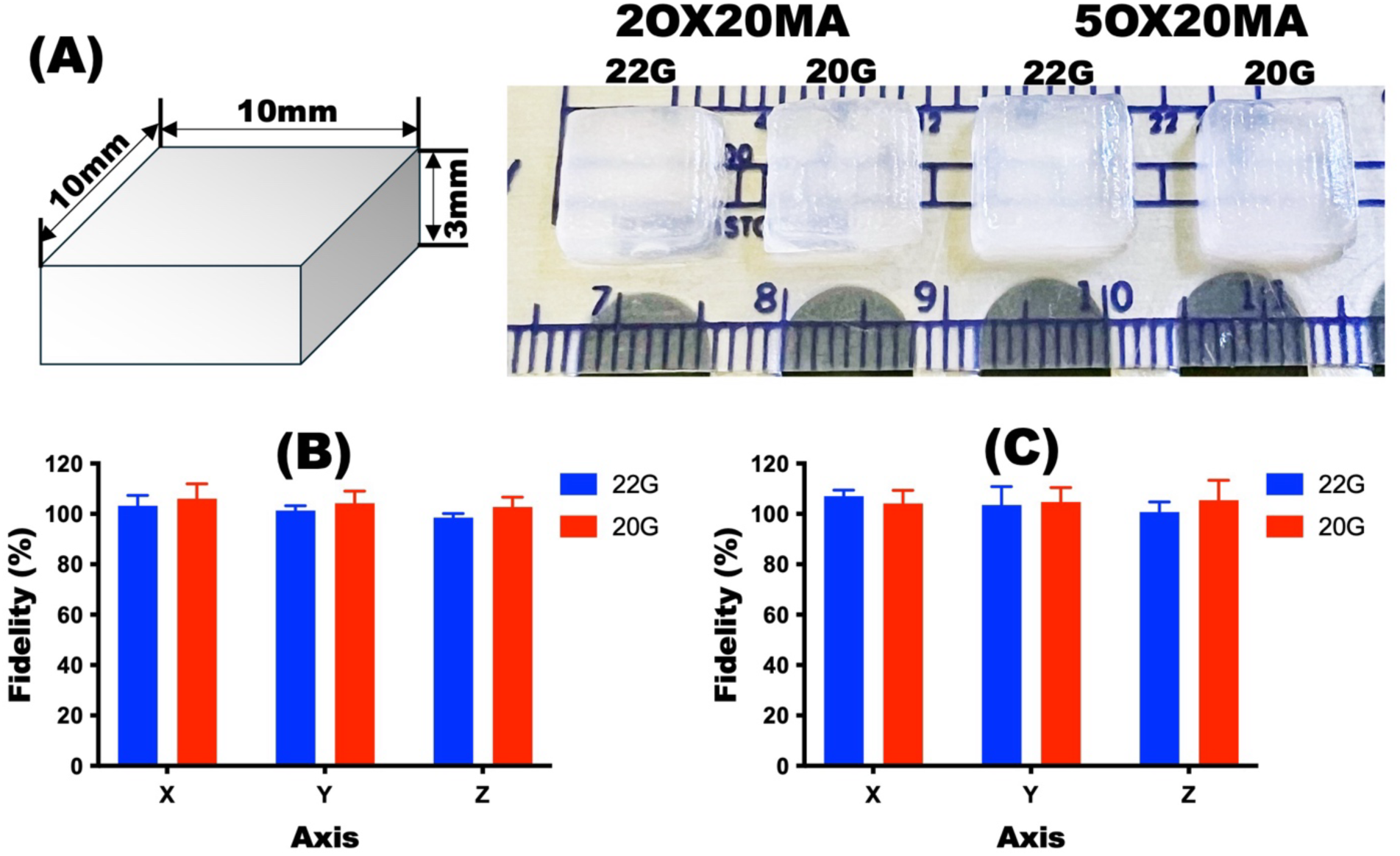
**(A)** Images of the photocrosslinked 3D printed structures using the hMSC encapsulated SSAM bioinks formed with 2OX20MA and 5OX20MA OMAs with 22G and 20G printing nozzles and quantified shape fidelity of **(B)** 2OX20MA and **(C)** 5OX20MA (N=3).

In native tissues, most cells are located within approximately 200μm of the nearest blood vessel, a spacing that allow for the effective diffusion of oxygen, and nutrient and waste removal to support tissue viability [39]. To develop clinically relevant tissue constructs that remain viable upon implantation, it is essential that the engineered tissue can establish microvascular networks [26]. Many studies have demonstrated that prevascularization of engineered constructs enhances vascularization and promotes functional tissue formation *in vivo* [24, 26, 40, 41]. Since the SSAM bioinks exhibit shear-thinning and self-healing properties and mechanical stability in the absence of shear forces, similar to materials reported in our previous studies [8, 10], the 3D bioprinted SSAM bioinks can also serve as slurry baths for 3D printing of individual cell-only bioinks. Therefore, prevascular network patterned constructs have been engineered by 3D printing of the SSAM bioink (**Figure 6A, i**) followed by bioprinting of individual HUVECs/hASCs prevascular bioink into the 3D printed microgel constructs (**Figure 6A, ii**) (**Supporting Movie 1**). Various prevascular network patterns have been successfully bioprinted into macro-scale tissue constructs (**Figure 6B**), and 3D bioprinted HUVECs/hASCs microfilaments exhibited high cell viability (**Supporting Figure 3**).

**Figure 6.**
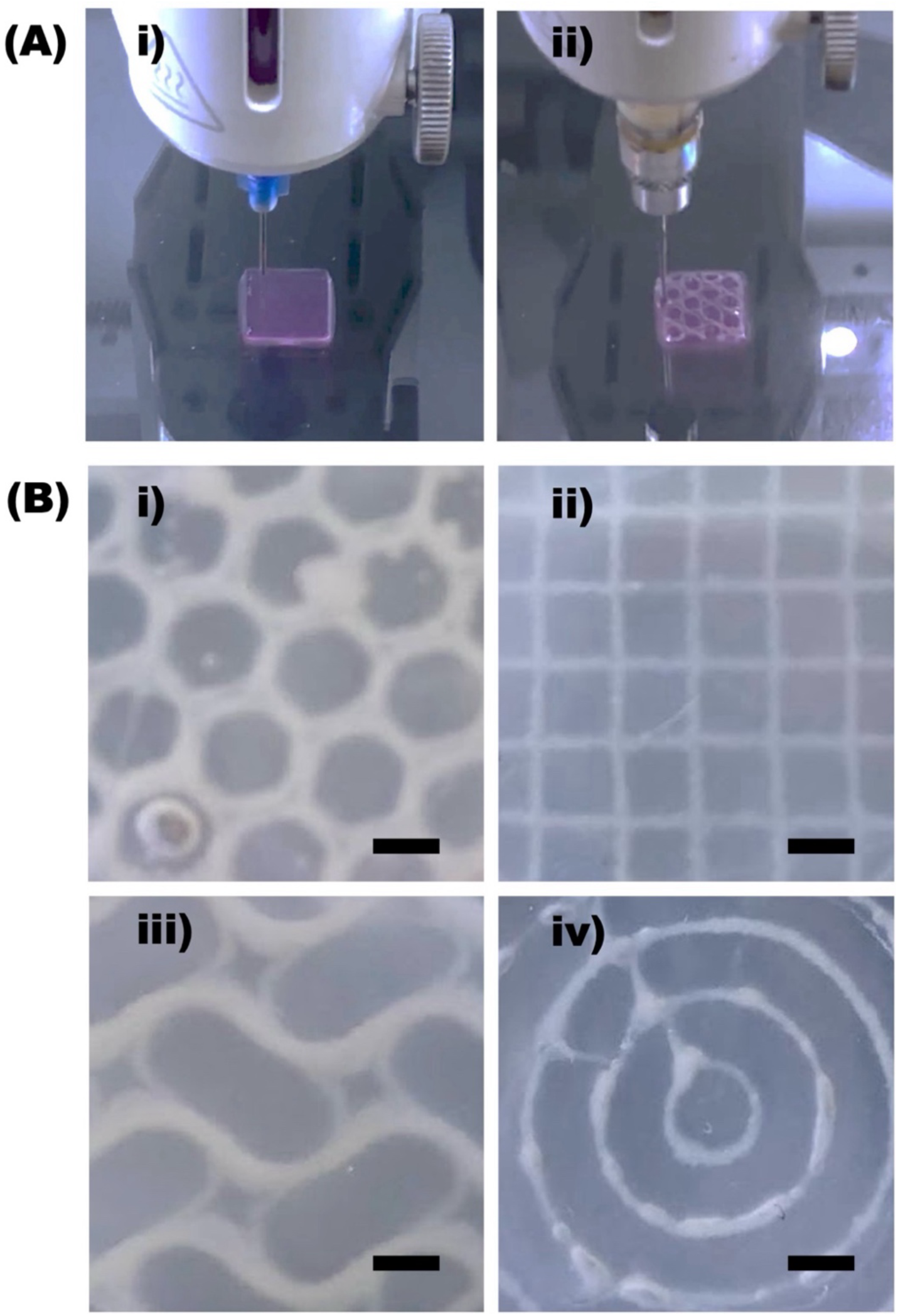
**(A)** Captured images during **i)** 3D printing of the SSAM bioink and **ii)** subsequent 3D printing of individual cell-only prevasculogenic bioink into the bioprinted 3D construct for prevascular network patterning. **(B)** Photographs of 3D prevascular networks formed into **i)** honeycomb, **ii)** grid, **iii)** wave and **iv)** concentric circles patterns by printing individual cell-only prevasculogenic bioinks into 3D slurries printed using the SSAM bioinks. Scale bars indicate 1 mm.

To demonstrate that the prevascularization can enhance functional tissue formation *in vitro*, prevascular network-patterned constructs have been fabricated by 3D bioprinting of hMSC-laden 2OX20MA or 5OX20MA SSAM bioinks followed by bioprinting of individual HUVECs/hASCs prevascular bioink into the 3D printed microgel constructs as described above. After 4 weeks of osteogenic differentiation, 3D printed prevasculature with grid pattern were well maintained in the 3D printed construct using 2OX20MA SSAM bioink (**Supporting Figure 4A**). In contrast, the 3D printed prevasculature pattern in the 3D printed construct using 5OX20MA SSAM bioink slurry migrated, contracted, and formed a large aggregate (**Supporting Figure 4B**) due to the weaker mechanical properties of the photocrosslinked 5OX20MA SSAM bioink (**Supporting Figure 2F**) compared to the photocrosslinked 2OX20MA SSAM bioink (**Figure 3F**). Although the grid-patterned prevasculature was well maintained in the 3D printed constructs formed by hMSC-laden 2OX20MA SSAM bioinks, prevasculogenesis was very limited due to mimimal interactions between the individual cell-only prevasculogenic bioink and the hMSC-laden OMA microgels (**Supporting Figure 4C and D**).

To promote interactions between the individual cell-only prevasculogenic bioinks and the SSAM bioinks, RGD-containing peptides were then incorporated into the 2OX20MA SSAM bioinks. After engieering the prevascular network-patterned constructs via 3D bioprinting using 22G (**Figure 7A**) and 20G (**Supporting Figure 5A**) printing needles as describe above, the 3D biprinted constructs were osteogenically differentiatied for 4 weeks. As a comparative group, prevascularized 3D printed constructs without a pattern were also prepared (**Figure 7B** and **Supporting Figure 5B**). CD31 staining enabled visualization of the development of prevascular network structures in the engieered constructs (**Figure 7C and E** and **Supporting Figure 5C and E**). Compared to the non-patterned prevascularization group (**Figure 7D and F** and **Supporting Figure 5D and F)**, the patterned prevascularization group (**Figure 7C and E** and **Supporting Figure 5C and E**) exhibted a significantly higher prevasculogenesis, as confirmed by increased branch numbers (**Figure 7G** and **Supporting Figure 5G**), junction numbers (**Figure 7H** and **Supporting Figure 5H**), triple point numbers (**Figure 7I** and **Supporting Figure 5I**) and quadruple point numbers (**Figure 7J** and **Supporting Figure 5J**), indicating enhanced hierarchical prevascular complexity. To further validate these findings at the molecular level, we quantified the mRNA expression of cadherin 5 (CDH5), also known as vascular endothelial cadherin, a critical gene in vascularization (**Figure 7K**). Consistent with prevascular structural complexity obsereved in the CD31 staining, CDH5 expression was significantly higher in the patterned prevascularized osteogenic constructs than in the non-patterned prevascularized osteogenic constructs. Since CDH5 is required for endothelial cell integrity, junction formation, angiogenesis, and maturation of blood vessels [42–44], this result further supports the histomorphometric finding of enhanced prevascular network formation in the patterned osteogenic constructs, highlighting the potential role of CDH5 in promoting prevascular complexity within enginered tissue constructs. To investigate the effect of patterned prevascularization on osteogenesis of encapsulated hMSCs, the osteogenically differentiated 3D printed constructs were evaluated for ALP activity, a key marker for osteogenic differentiation of stem cells. The ALP activity of hMSCs in the prevascularization groups was significantly higher than in the no prevascularization group (**Figure 8A**), while only the ALP/DNA of hMSCs in the patterned prevascularization group was significantly higher than that of the no prevascularization group (**Figure 8B**). Since matrix mineralization in engineered constructs is the most important indicator of osteogenic differentiation, calcium depositon in the 3D printed constructs was also evaluated using quantification of calcium content and Alizarin red S staining. As shown in **Figure 8C and D**, calcium deposition in the prevascularization groups was significantly higher than in the no vascularization group. Consistent with the ALP activity results, mineralization was further enhaced by patterned 3D printing of the individual cell-only prevasculogenic bioinks. Similar to the calcium content results, the prevascularized constructs (**Figure 8E and F**) stained more intensely with Alizarin red S compared to the no prevascularization construct (**Figure 8G**), and the black regions in the patterned prevascularization constructs (**Figure 8E**) indicated the areas of strongest Alizarin red S staining, suggesting that patterned prevascularization further intensified mineral depostion in the constructs. The osteogenically differentiated 3D printed constructs using 20G printing needles (**Supporting Figure 6**) exhibited similar trends compared to constructs printed with 22G printing needles. The expression of osteocalcin (OCN), an important marker in the mineralization process during osteogenic differentiation of hMSCs, was significantly increased in the prevascularized osteogenic constructs and further enhance by patterned prevascularization (**Supporting Figure 7**), consistent with the higher levels of mineralization. These findings highlight the potential of prevascular patterning in promoting osteogenesis of hMSCs within engineered tissue constructs, suggesting that structured prevascular networks may provide a supportive environment for bone tissue formation. These results underscore the value of incorporating defined prevascular designs in engineered constructs, potentially advancing stategies in bone tissue engineering and other regenerative medicine applications. Moreover, these results suggest that future work should focus on optimizing prevascular patternig, exploring the interactions between individual cell-only bioinks and stem cell-laden microgel bioinks, and developing scaling methods to facilitate the clinical translation of these findings.

**Figure 7.**
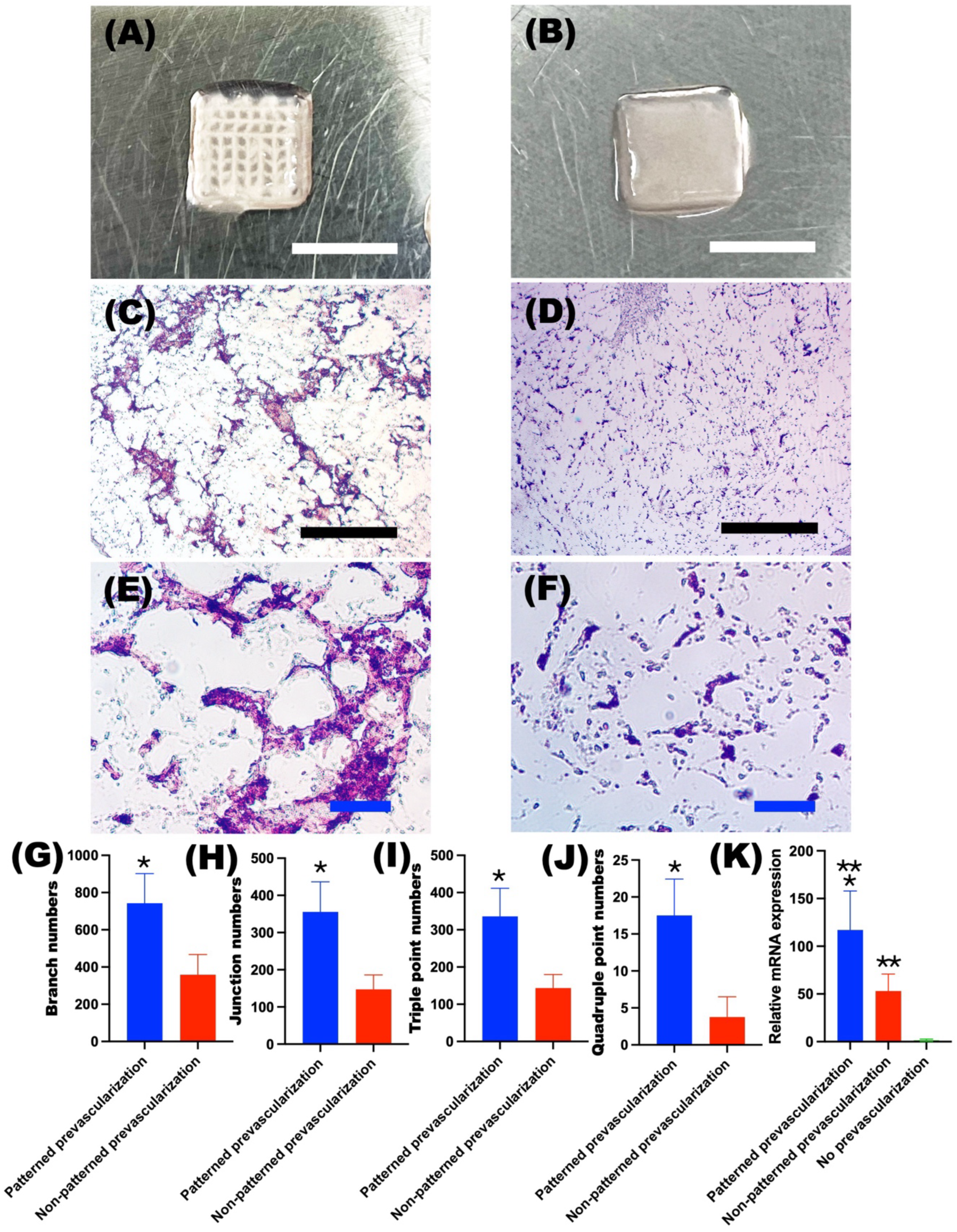
Photographs of **(A)** 3D printed patterned prevascularization and **(B)** non-patterned prevascularization in constructs. Photomicrographs of CD31 stained **(C)** 3D printed patterned prevascularization and **(D)** non-patterned prevascularization in constructs at low magnification, and **(E-F)** their high magnification images. Quantified prevasculogenesis by measuring **(G)** branch numbers, **(H)** junction numbers, **(I)** triple point numbers, and **(J)** quadruple point numbers using AnalyzeSkeleton program from ImageJ (N=4). (**K**) Relative CDH5 gene expression in the osteogenically differentiated 3D printed constructs (N=4). The relative gene expression levels were normalized using the no prevascularization group. *\*p*<0.05 compared to the non-patterned prevascularization group. *\*\*p*<0.05 compared to the no prevascularization group.

**Figure 8.**
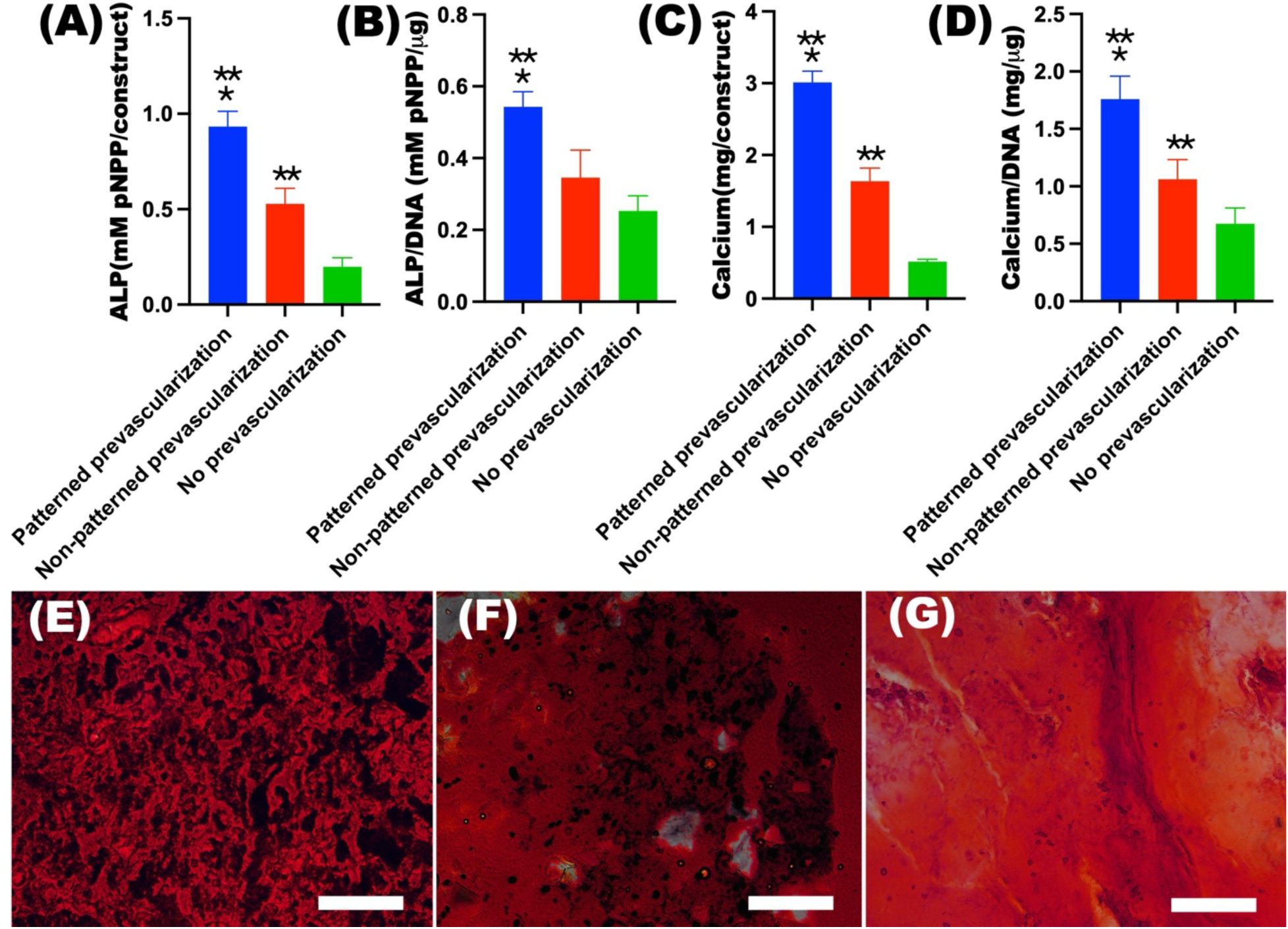
Quantification **(A)** ALP, **(B)** ALP/DNA, (**C)** calcium, and **(D)** calcium/DNA in the 3D printed constructs (using a 22G needle) after 4 weeks culture (N=4). Photomicrographs of Alizarin red S stained **(E)** 3D printed patterned prevascularization, **(F)** non-patterned prevascularization and **(G)** no prevascularization in constructs after 4 weeks culture. *\*p*<0.05 compared to the non-patterned prevascularization group. *\*\*p*<0.05 compared to the no prevascularization group.

*In situ* bioprinting is an emerging technology that enables the direct printing of bioinks into a patient’s body during surgery, allowing for precise placement of printed materials at specific locations to promote the seamless integration of engineered tissues with native host tissues. By eliminating the costly and time intensive *in vitro* culture process, *in situ* bioprinting offers promising solutions for tissue engineering and regeneration, especially in complex wounds or defect sites. However, *in situ* bioprinting faces challenges, such as the difficulty of developing optimal bioinks that are biocompatible and posses the necessary mechanical properties for precise deposition within the patient’s body during surgery. Additionally, ensuring bioink printability, mechanical stability under physiological conditions after printing and maintenance of sterility throughout the process are important factors to consider. We demonstrate here that the SSAM bioinks can overcome these challenges by *in situ* 3D bioprinting SSAM bioink into a mouse calvarial bone defect followed by 3D printing of indiviual cell-only bioink into the 3D bioprinted SSAM construct (**Figure 9A and B** and **Supporting Movie 2**). After photocrosslinking (**Figure 9C**), the mechanically stable prevascularized construct was successfully extracted from the calvarial defect (**Figure 9D**). This result highlights the potential and feasibility of our approach for clinical applications, as our SSAM bioinks and individual cell printing technique enable *in situ* bioprinting of prevascularized tissue constructs by achieving mechanical stability, biocompatibility and integration with host tissues.

**Figure 9.**
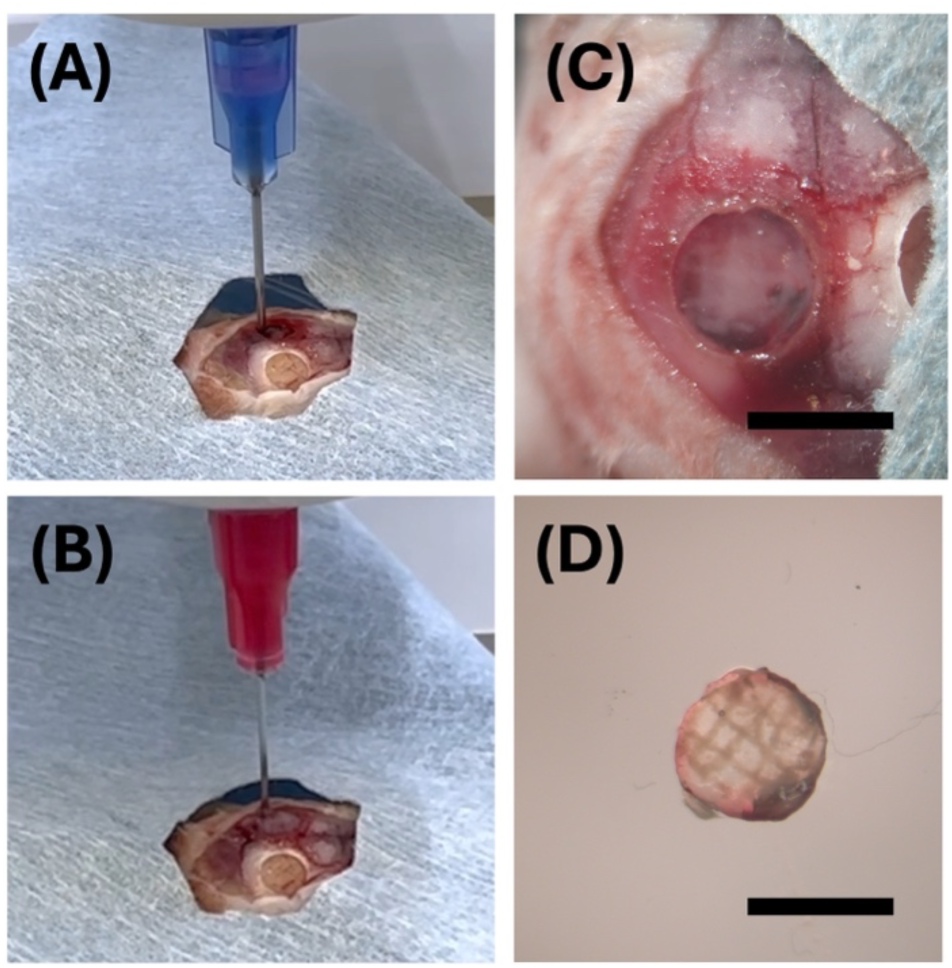
Captured images during **(A)** *in situ* 3D printing of the SSAM bioink into a mouse calvarial bone defect and **(B)** subsequent 3D printing of individual cell-only prevasculogenic bioink within the bioprinted 3D construct. **(C)** Photographs of *in situ* 3D bioprinted prevascular network patterned construct after photocrosslinking and **(D)** extracted construct from the calvarial bone defect. The scale bars indicate 4 mm.

## Conclusions

The SSAM bioinks developed in this study demonstrate the potential for advancing 3D bioprinting in tissue engineering and regeneration medicine applications, particularly for engineering vascularized tissue constructs. The SSAM bioinks exhibit shear-thinning, low shear yield stress, and self-healing properties with high cell viability, which are crucial for achieving high resolution printing and the fabrication of complex 3D structures with high shape fidelity. Moreover, the incorporation of prevascular networks using an individual cell-only printing technique promotes vascularization and enhances osteogenic differentiation of the 3D printed tissue constructs, providing a promising platform for bone tissue engineering applications. This study highlights the feasibility of *in situ* 3D bioprinting for clinical applications, paving the way for more effective, personalized and functional tissue regeneration.

## Acknowledgements

The authors gratefully acknowledge funding support from the Department of Veterans Affairs, Veterans Health Administration, Office of Research and Development, Rehabilitation Research and Development Service under award number I01RX004825 and the National Institutes of Health’s National Institute of Arthritis and Musculoskeletal and Skin Diseases under award number R01AR081448. The contents of this publication are solely the responsibility of the authors and do not necessarily represent the official views of the Department of Veterans Affairs or the National Institutes of Health.

## Notes

### Competing Interest Statement

The authors have declared no competing interest.

