## Supplemental figures and table for "*In situ* cell-only bioprinting of patterned prevascular tissue into bioprinted high-density stem cell-laden microgel bioinks for vascularized bone tissue regeneration"

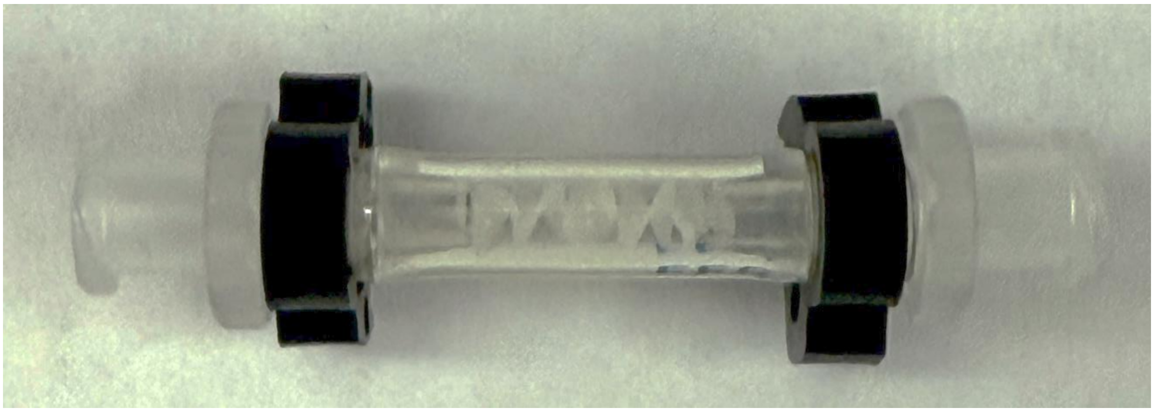

**Supporting Figure 1.** Photograph of the custom female-female lure lock spiral mixing unit.

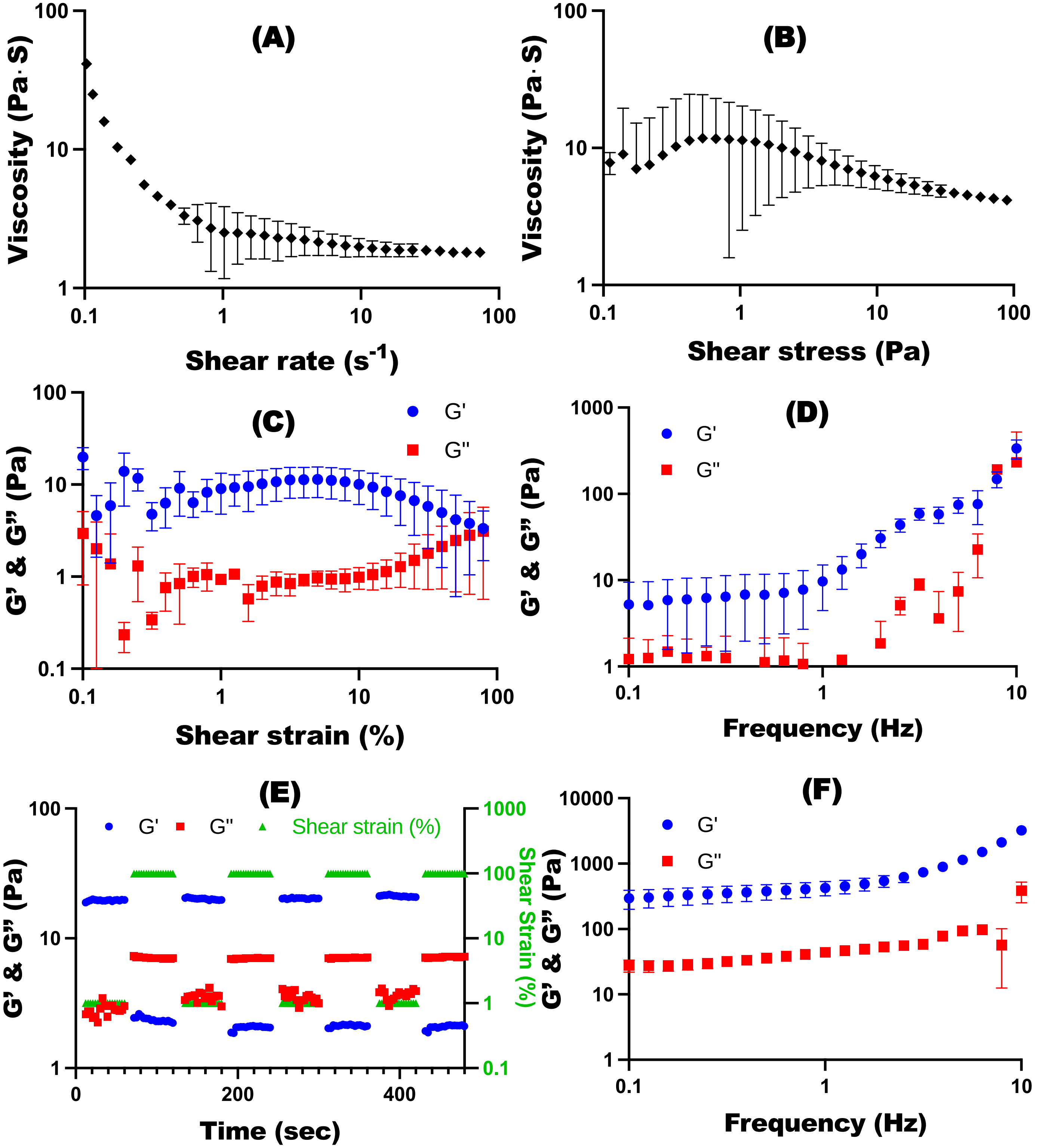


**Supporting Figure 2.** Viscosity measurements of the SSAM bioinks fabricated with 5OX20MA OMA as a function of **(A)** shear rate and **(B)** shear stress demonstrate their shear-thinning and shear-yielding behaviors (N=3). **(C)** Frequency sweep tests indicate that the SSAM bioinks were mechanically stable (N=3). **(D)** Strain sweep tests of the SSAM bioinks indicate their gel-sol transition at approximately 90 % shear strain (G' and G" crossover) (N=3). **(E)** Shear moduli changes during dynamic shear strain test of the SSAM bioinks with alternating low (1%) and high (100%) shear strains at 1Hz frequency demonstrate their self-healing or thixotropic property by showing rapid transition between solid-like and liquid-like behaviors. **(F)** Frequency sweep tests of the photocrosslinked hMSC encapsulated SSAM bioinks indicate that additional photocrosslinking enhanced their mechanical strength (N=3).

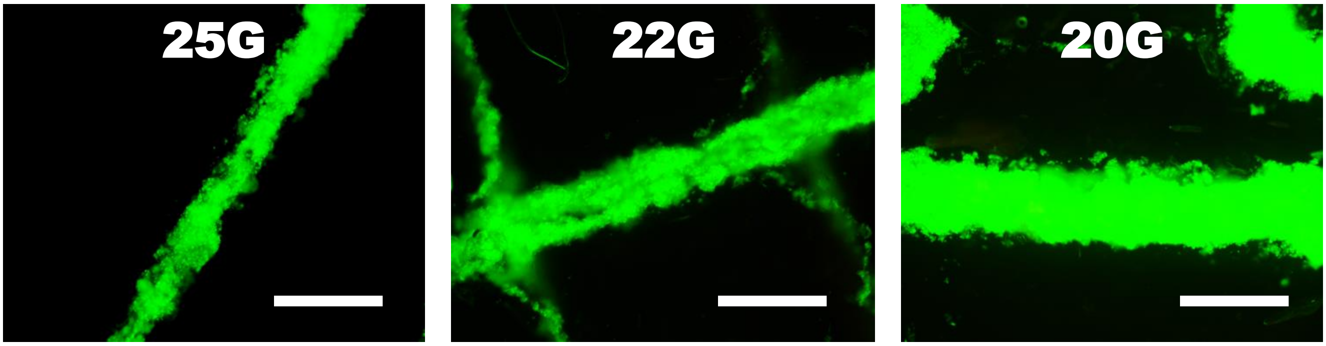


**Supporting Figure 3.** Representative Live(green)/Dead(red) photomicrographs of 3D printed HUVECs/hASCs microfilaments in the 2OX20MA SSAM bioinks using various printing needles. Scale bars indicate 500 μm.

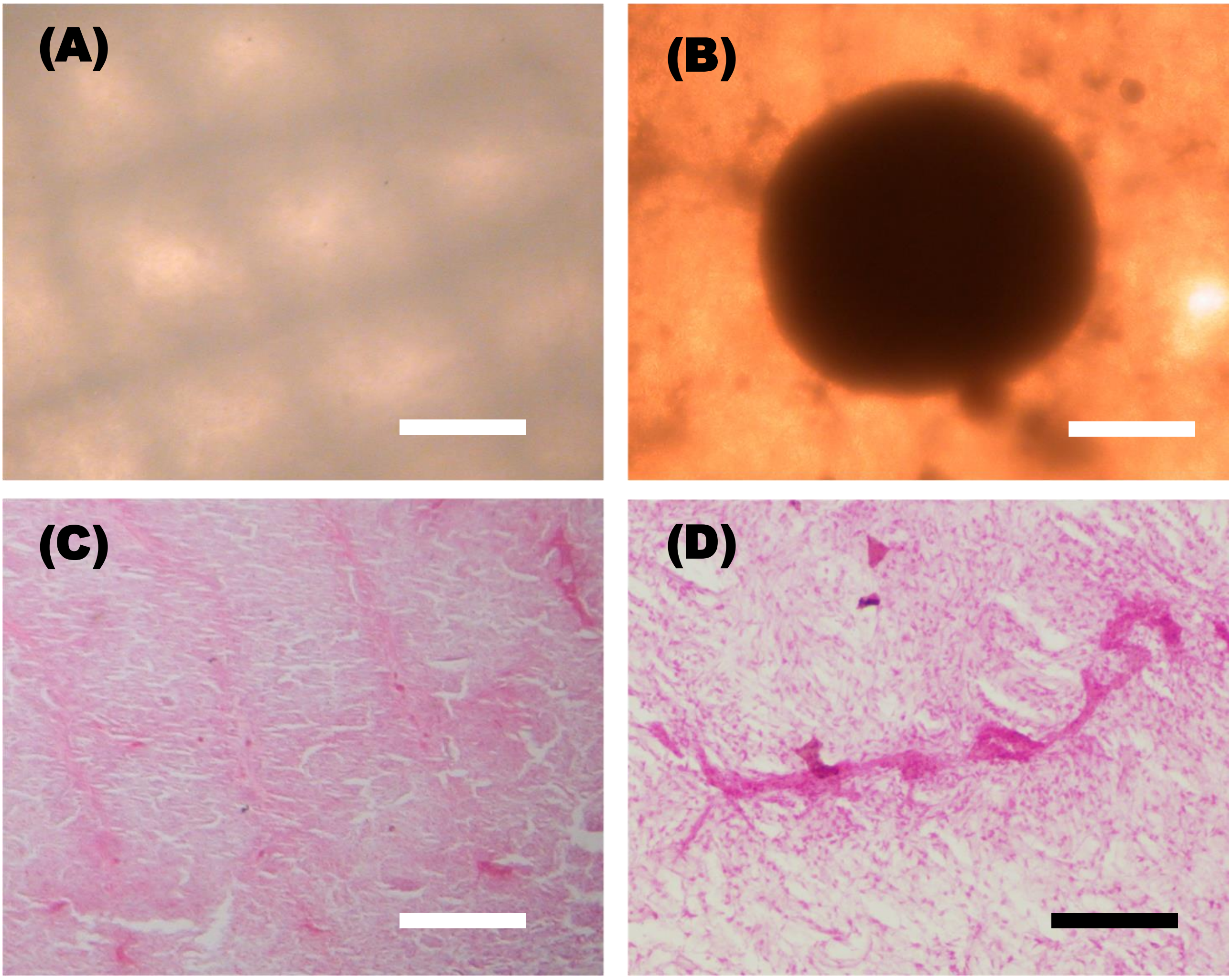


**Supporting Figure 4.** Photomicrographs of osteogenically differentiated 3D printed prevascular network-patterned constructs using **(A)** 2OX20MA and **(B)** 5OX20MA SSAM bioinks. Photomicrographs of CD31 stained osteogenically differentiated 3D printed prevascular network-patterned constructs using 2OX20MA SSAM bioinks at **(C)** low and **(D)** high magnification. White scale bars indicate 1mm. Black scale bar indicates 200 µm.

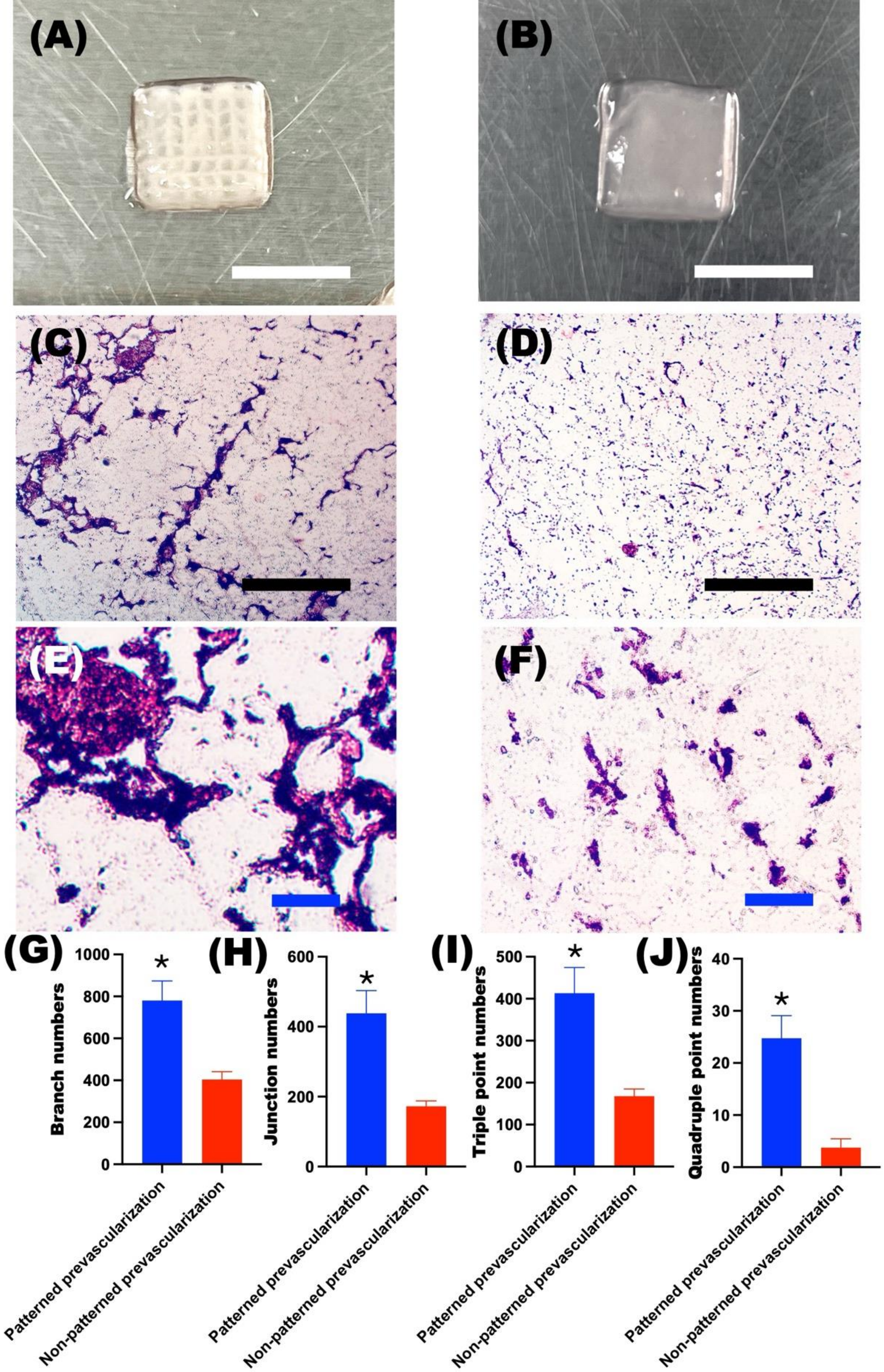


**Supporting Figure 5.** Photographs of **(A)** 3D printed patterned prevascularization and **(B)** non-patterned prevascularization in constructs. Photomicrographs of CD31 stained **(C)** 3D printed patterned prevascularization and **(D)** non-patterned prevascularization in constructs at low magnification, and **(E-F)** their high magnification images. Quantified prevasculogenesis by measuring **(G)** branch numbers, **(H)** junction numbers, **(I)** triple point numbers, and **(J)** quadruple point numbers using AnalyzeSkeleton program from ImageJ. *$p<0.05$ compared to the non-patterned prevascularization group (N=4).

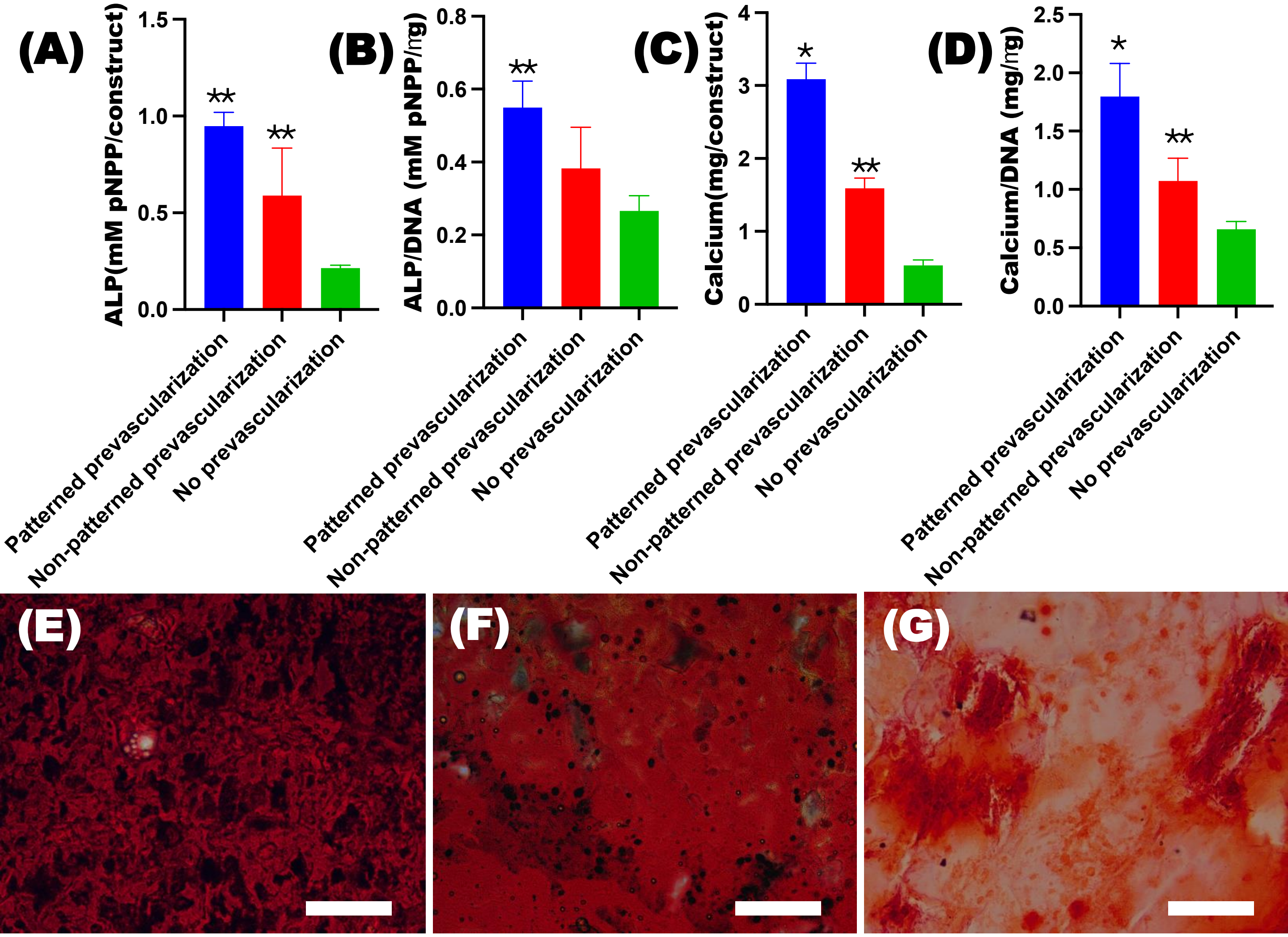


**Supporting Figure 6.** Quantification **(A)** ALP, **(B)** ALP/DNA, (**C)** calcium, and **(D)** calcium/DNA in the 3D printed constructs using a 20G needle after 4 weeks culture (N=4). Photomicrographs of Alizarin red S stained **(E)** 3D printed patterned prevascularization, **(F)** non-patterned prevascularization and **(G)** no prevascularization in constructs after 4 weeks culture. . $*p<0.05$ compared to the non-patterned prevascularization group. $**p<0.05$ compared to the no prevascularization group.

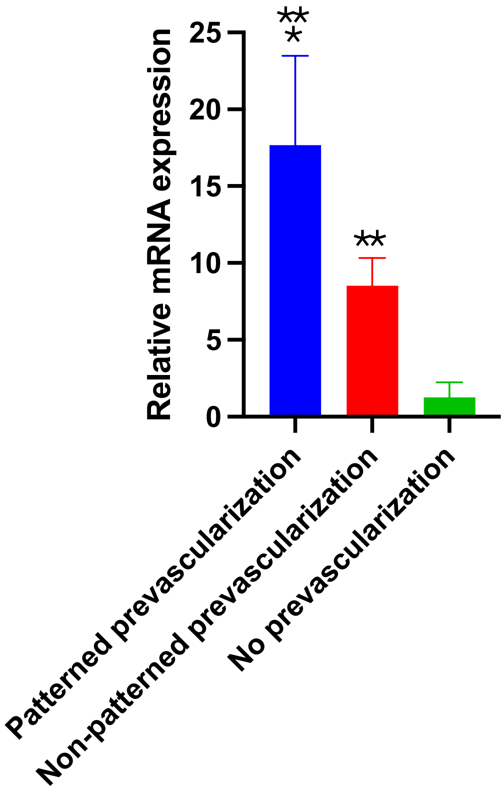


**Supporting Figure 7.** Relative OCN gene expression in the osteogenically differentiated 3D printed constructs. The relative gene expression levels were normalized using the no prevascularization group (N=4). *$p$<0.05 compared to the non-patterned prevascularization group. **$p$<0.05 compared to the no prevascularization group.

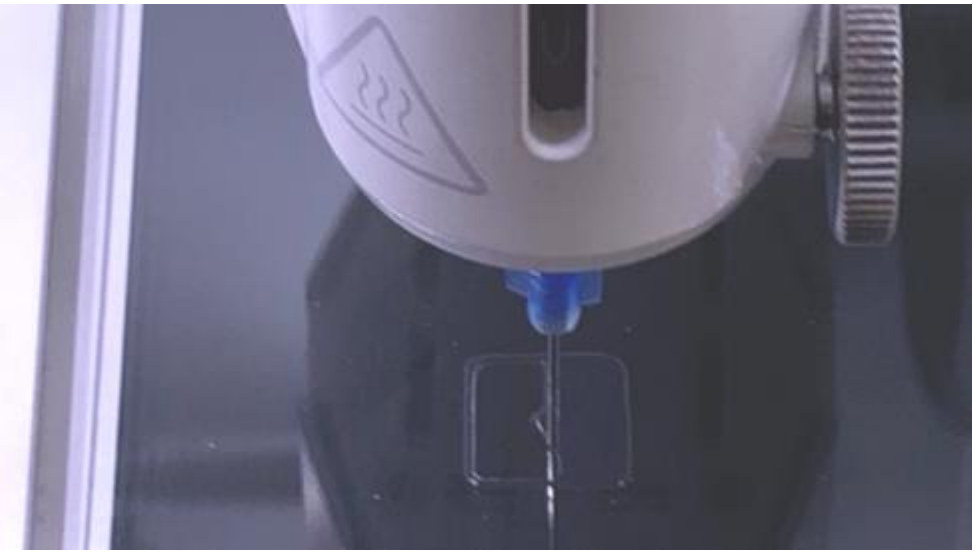

**Supporting Movie 1.** 3D printing of the SSAM bioink and subsequent 3D printing of individual cell-only prevasculogenic bioink into the bioprinted 3D construct for prevascular network patterning.

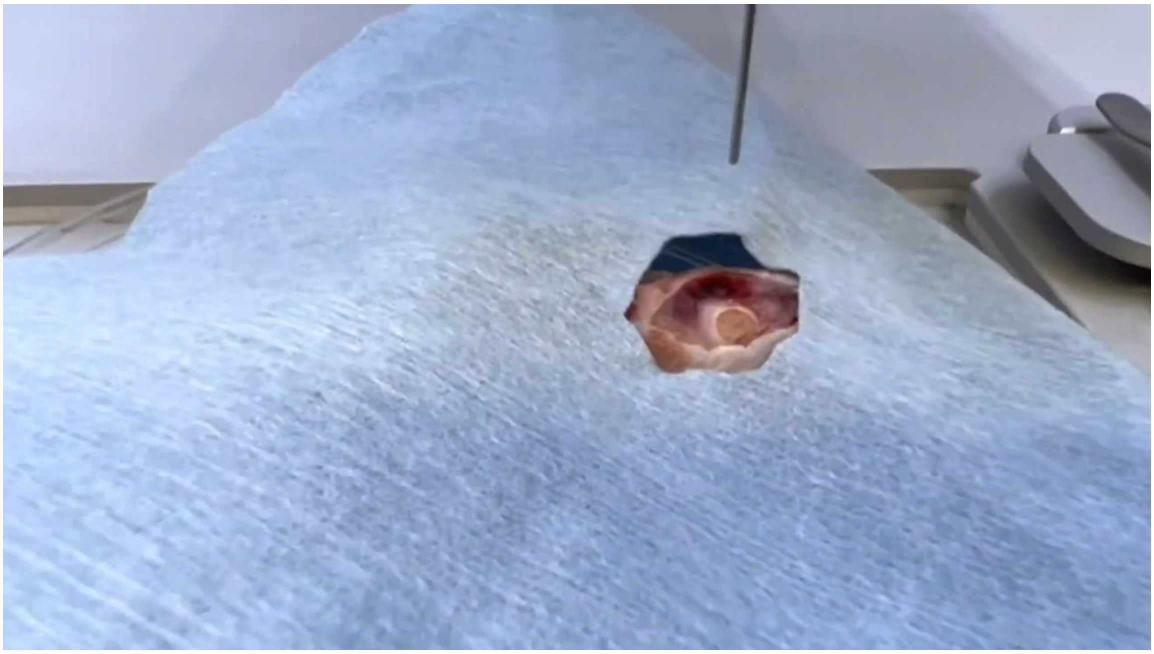

**Supporting Movie 2.** *In situ* 3D printing of the SSAM bioink into a mouse calvarial bone defect and subsequent 3D printing of individual cell-only prevasculogenic bioink into the bioprinted 3D construct.

**Supporting Table 1.** Oligonucleotide primer sequences for qRT-PCR.

| Gene | Sequence (5'-3') | | Accession number |
|---|---|---|---|
| GAPDH | Forward | GGGGCTGGCATTGCCCTCAA | NM_002046 |
| | Reverse | GGCTGGTGGTCCAGGGGTCT | |
| CDH5 | Forward | GAAGCCTCTGATTGGCACAGTG | NM_001795 |
| | Reverse | TTTTGTGACTCGGAAGAACTGGC | |
| OCN | Forward | ATGAGAGCCCTCACACTCCTC | NM_199173 |
| | Reverse | CGTAGAAGCGCCGATAGGC | |
